# Connexin36 expression in the cochlear nucleus complex of the echolocating bat, *Eptesicus fuscus*

**DOI:** 10.1101/2022.03.23.485527

**Authors:** Alyssa W. Accomando, Mark A. Johnson, Madeline A. McLaughlin, James A. Simmons, Andrea Megela Simmons

**Author notes:** Present address: Taconic Biosciences, Rensselaer, NY 12144, USA.

## Abstract

Gap junctions and electrical synapses in the central nervous system are associated with rapid temporal processing and coincidence detection. Using histology, immunohistochemistry, and *in situ* hybridization, we investigated the distribution of Connexin36 (Cx36), a protein that comprises neuronal gap junctions, throughout the cochlear nucleus complex of the echolocating big brown bat, *Eptesicus fuscus*, a species exhibiting extreme behavioral sensitivity to minute temporal changes in ultrasonic echoes. For comparison, we visualized Cx36 expression in the cochlear nucleus of transgenic Cx36 reporter mice, species that hear ultrasound but do not echolocate. We observed Cx36 expression in the anteroventral and dorsal cochlear nucleus, with more limited expression in the posteroventral cochlear nucleus, of both species. Several different morphological cell types were labeled, including globular and spherical bushy, octopus, stellate, and fusiform cells. Labeled Cx36 puncta were also observed. Cx36 expression in the bat was spread throughout a relatively smaller area of the cochlear nucleus than in the mouse, even though the bat cochlear nucleus is hypertrophied. In the bat, the anteroventral cochlear nucleus showed higher percent area label than the dorsal cochlear nucleus, with a trend towards the opposite result in the mouse. The presence of gap junctions appears to be a conserved feature of the mammalian cochlear nucleus and thus not uniquely tied to the temporal hyperacuity of echolocation.

## INTRODUCTION

Gap junctions between neurons mediate transmission through electrical synapses. They are important features of brain function because they co-exist with chemical synapses, allow for bi-directional electrical coupling between cells, are prominent in inhibitory interneurons, and exhibit activity-dependent plasticity (Bennett and Zukin 2004; Pereda 2014; Connors 2017). Electrical coupling can coordinate activity in groups of neurons without the reduced timing accuracy of chemical synapses (Pereda 2014). The role of gap junctions in synchronizing neuronal activity has inspired a search for electrical synapses in the auditory system, where precise timing and coincidence detection are crucial for periodicity (pitch) perception and the localization of sounds in space (Joris et al. 1998; Simmons and Simmons 2011; Galaretta and Hestrin 2001, Finneran et al. 2020). The hypothesized presence of neuronal circuits specialized for accurate registration of the timing of neighboring sound frequencies is the signature feature of computational auditory models (Lyon 2017).

The mammalian cochlear nucleus (CN) receives direct synaptic input from the auditory (8th) cranial nerve. Bushy cells in the ventral cochlear nucleus [VCN, comprising the anteroventral (AVCN) and the anterior portion of the posteroventral (PVCN) cochlear nuclei] exhibit more precise synchronization to the envelope of sounds than do auditory nerve fibers, and convey this temporal information to other brainstem auditory nuclei (Joris and Smith 2008; Rubio 2018). Cells in the dorsal cochlear nucleus (DCN) receive auditory (VCN, auditory nerve) and multimodal input, providing information for localization of sounds in elevation and for head and ear position (Trussell and Oertel 2018). The presence of gap junctions has been confirmed electrophysiologically in the CN of several mammalian species (rat: Sotelo et al. 1976; Wouterlood et al. 1984; Mugnaini 1985; Gomez-Nieto and Rubio 2009; monkey: Gomez-Nieto and Rubio 2011; mouse: Apostolides and Trussell 2013, 2014a,b; Yaeger and Trussell 2016). Because gap junctions are composed of connexin proteins, connexins serve as immunohistochemical markers for the presence of electrical synapses (Rash et al. 2000). Connexin36 (Cx36) is the main neuronal connexin in the adult mammalian brain and spinal cord (Condorelli et al. 1998; Belluardo et al. 2000; Rash et al. 2000). In mice and rats, Cx36 labels cells in the DCN, AVCN, medial nucleus of the trapezoid body, lateral superior olivary nucleus, dorsal nucleus of the medial lemniscus, inferior colliculus, and auditory cortex (Rubio and Nagy 2015). In goldfish, Cx35 (the fish ortholog for mammalian Cx36) is expressed in Mauthner cells, which receive mixed electrochemical auditory nerve afferents (Pereda et al. 2003) and mediate the fast tail flip escape/startle response (Eaton et al. 1977).

Insectivorous bats use echolocation for navigation and prey catching (Griffin 1958). Bats emitting frequency-modulated echolocation calls (FM bats) perceive the distance to a target, such as a small insect, by measuring the time delay between each emitted call and the returning echo; they classify the size and shape of the target by the echo’s internal temporal and spectral structure (Simmons et al. 1995, 2014; Yovel et al. 2011). Big brown bats (*Eptesicus fuscus*) exhibit extreme sensitivity for detecting microsecond or even smaller changes in echo delay and phase (Simmons et al. 2003; Bates and Simmons 2011; Simmons 2014). This acute timing perception has important technological implications (Balieri et al. 2017), prompting efforts to identify neurobiological underpinnings and develop computational models of FM echolocation (Covey and Casseday 1995, 1999; Sanderson et al. 2003; Covey 2005; Simmons 2012, 2014; Faure and Firzlaff 2016; Luo et al. 2018; Ming et al. 2021).

An earlier report from our laboratory used immunohistochemistry to describe Cx36 expression, and the possible presence of gap junctions, in the CN of the big brown bat (Horowitz et al. 2008). Results showed label in dorsal regions of the AVCN and DCN, with reduced expression in the PVCN. In comparison experiments using identical techniques, no label was seen in the mouse CN. Given these apparent species differences, the authors speculated that electrical neurotransmission in the bat’s CN might be a neurobiological specialization for the fine temporal acuity of echolocation. Since then, Cx36 immunohistochemical label has been shown in the CN of adult mice (C57 BL/6 and CD1; Rubio and Nagy 2015), raising questions about the methodology used by Horowitz et al. (2008). The present study re-examined Cx36 expression in the CN complex of the big brown bat, this time using multiple techniques including immunochemistry and *in situ* hybridization, and compared these results to Cx36 label in the CN of transgenic reporter mice. These experiments shed light on the role of electrical synapses in early-stage auditory processing by mammals, but also address their hypothesized unique role in echolocation.

## METHODS

Adult big brown bats (*n* = 7; ages unknown) were wild caught under a State of Rhode Island scientific collection permit, which limits the number of animals that can be caught during a restricted seasonal period. They were housed in a Level 2 biohazard space on a 12-hour reverse light-dark cycle, maintained at approximately 23° C and 60-75% humidity to prevent torpor. Bats were fed mealworms in quantities allowing them to maintain healthy body weights of 15-18 g. They had access to vitamin-enriched water *ad libitum*. Adult (6 months old) male mice heterozygous for Cx36/LacZ-IRES-PLAP on a C57BL/6 background (n=4) and wild-type littermates (n=2) were obtained from the laboratory of Gilead Barnea in the Neuroscience Department of Brown University. All procedures were approved by the Brown University Institutional Animal Care and Use Committee and adhered to federal regulations for animal experimentation.

### Cytoarchitecture

Bats (*n* = 2) were terminally anesthetized by intraperitoneal injection of Beuthanasia-D (active ingredients: pentobarbital sodium and phenytoin). After cessation of all reflexes, they were transcardially perfused with 0.9% heparinized saline followed by 4% paraformaldehyde (PFA) in phosphate buffered saline (PBS, pH 7.4). The head of one bat was removed and placed in decalcifying solution (Richard Allan Scientific) for 3 weeks, with fresh solution every 5–7 days. Following decalcification, the head was stripped of external epidermis, embedded in paraffin, and sectioned coronally at 5 μm thickness on a cryostat (Leica 3050). Sections were mounted to glass slides and stained with trichrome. The brain of the second bat was removed from the head, embedded in 5% agarose (ISC Bioexpress) in PBS (pH 7.4), then sectioned coronally on a vibratome at 40 μm thickness and stained with 0.5% cresyl violet acetate. Brightfield microscopy was performed on an Olympus BX60 microscope (Olympus Scientific); images were acquired using an Olympus DP72 camera and cellSens software, which was used to white-balance images.

For standardization with previous anatomical work and to contextualize the results from this study, boundaries of the CN subnuclei were identified from an existing atlas of the *E. fuscus* brain (Carter et al., 2004), based on cresyl violet histology of the brain of one adult female bat. To obtain the two-dimensional area of CN subnuclei, individual images (n = 8 for AVCN, n = 7 for PVCN, and n = 5 for DCN) from this atlas were measured within the prescribed boundaries by an experimenter blinded to the hypothesis using the manual Freehand tool in ImageJ. These 2D measurements were then multiplied by section thickness (50 μm) to obtain volume; and, since there was 150 μm between sections, the average area of sequential images was multiplied by 150 to estimate the volume between samples. Volumes of all images and the estimated volumes between samples were added to obtain final volume estimates. No corrections were applied to account for 9.2% estimated tissue shrinkage (Carter et al., 2004).

### Immunohistochemistry

Bats (*n* = 3) were terminally anesthetized and transcardially perfused as described above. Heads were removed, excess tissue was dissected away, and samples were preserved in 4% PFA overnight at 4° C. Brains were then dissected and post-fixed overnight. They were then cryoprotected in 30% sucrose (w/v) in distilled water and embedded in OCT (Tissue-Tek, Sakura Finetek) on dry ice. Tissue was sectioned at 20 μm on a cryostat, divided into four sets, and mounted on charged slides (Fisherbrand Superfrost Plus, Thermo Fisher Scientific). Sections were stored in airtight containers at −80°C until further processing. Before immunohistochemistry (IHC) was performed, brain tissue was thawed in a humidified chamber for 30 min to 1 hr. Sections were then postfixed (4% PFA) for 20 min, washed in PBS, and subjected to antigen retrieval with sodium citrate buffer (10 mM tri-sodium citrate dihydrate, 0.05% Tween 20, pH 6.0) at 80° C for 20 min. Different sets of sections were processed for diamobenzidine (IHC-DAB) and immunofluorescence (IHC-F).

#### IHC-DAB

Although the antigen retrieval step quenches endogenous peroxidases, sections were further quenched under a Parafilm coverslip in 3% H_2_O_2_ in PBS for 1 hr at room temperature inside a humidified chamber. Three 10 min washes in PBST (phosphate-buffered saline with 0.05% Tween-20) were performed, then hydrophobic barriers were placed around the edge of each glass slide such that the remainder of processing steps could be performed directly on each slide. Protein binding sites were blocked using bovine serum albumin (BSA) and normal goat serum (NGS) (in PBS with 0.006% triton) for 1 hr at room temperature. Because brain tissue contains endogenous biotin, streptavidin and biotin blocking steps (15 min each under Parafilm coverslips in a humidified chamber) were performed. Sections were then incubated in primary antibody in PBS with 0.5% BSA overnight at 4° C in a humidified chamber (see Table 1 for all antibody manufacturers and dilutions). Sections were washed in PBST and incubated in an HRP-conjugated secondary antibody for 2 hr at room temperature. After washing, a tyramide-biotin, streptavidin-HRP amplification step was performed. Two controls were added: (1) omitting primary antibody and (2) omitting the tyramide-biotin amplification step. The DAB reaction was performed to produce a brown-colored reaction product. A Hematoxylin counterstain was performed and blued using Scott’s tap water (20 g/L magnesium sulfate and 2 g/L sodium bicarbonate). Sections were washed with distilled water, cover slipped with AquaMount (Fisher Scientific), and stored at room temperature. Brightfield microscopy was performed on an Olympus BX60 microscope (Olympus Scientific); images were acquired using an Olympus DP72 camera and cellSens software, which was used to white-balance images. Images were then analyzed to count and measure the area of DAB-labeled cells using the manual Freehand tool in ImageJ by an experimenter blinded to the hypothesis.

**Table I.**
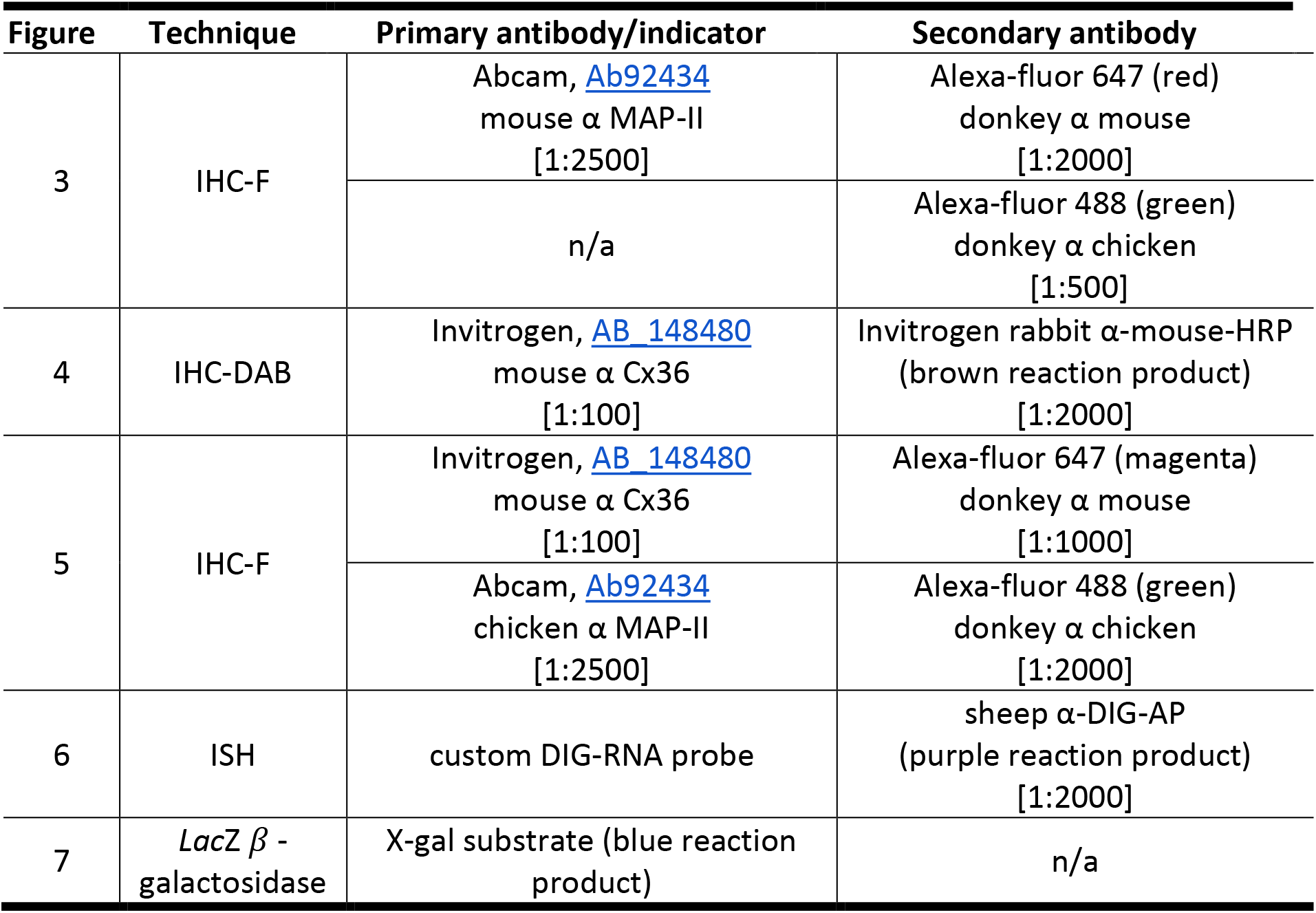
List of figures and assay techniques, including links to The Antibody Registry.

#### IHC-F

IHC-F was performed on different sets of sections from the three bat brains described above, as well as from a fourth brain not used for IHC-DAB. Thawed cryosections were subjected to post-fixation, washes, sodium-citrate antigen retrieval, blocking steps, and overnight primary antibody incubation at 4° C (Table 1). Fluorophore-conjugated secondary antibodies (Abcam) were applied for 2 hr at room temperature. Two sets of sections (one set from each of two brains) were labelled with a polyclonal antibody to microtubule-associated protein II (MAP-II), a neuron-specific marker. In one set, a high concentration [1:100] of secondary antibody was also applied in order to non-specifically label the background tissue. In another set, an additional monoclonal primary antibody against Cx36 was used to co-label neurons with Cx36. Finally, slides were washed and mounted with ProLong Gold Antifade Mountant with DAPI (Fisher Scientific) and stored at room temperature. Three-dimensional image stacks (z-stacks) were acquired from fixed samples using a Zeiss LSM 510 confocal laser scanning microscope. Fluorophores were excited by lasers of different wavelengths [blue (DAPI, 460 nm), green (Fluorescein, 535 nm), and red (Rhodamine, 620 nm)], then images were optimized and captured using Zeiss software. Collapsed z-stack images were then analyzed to count and measure the area of both labeled neurons and Cx36 puncta by an experimenter blinded to the hypothesis using the automated Analyze particles tool in ImageJ.

### Cx36 in situ hybridization

#### Bat

An *in-situ* hybridization (ISH) antisense probe was generated from big brown bat genomic DNA (Fig. 1). The Cx36 protein is encoded by the GJD2 gene, which was identified in the big brown bat using the known mouse sequence [MGI:1334209 (Condorelli et al., 1998)]. The sequences of connexin genes are highly conserved across mammalian phylogeny, so a tBLASTn search was performed against big brown bat genome sequence data [BioProject PRJNA72449, Di Palma et al. (2012)], and a 99% match was found. Based on this newly identified big brown bat ortholog sequence, oligonucleotide primers for generating an ISH template (IDT DNA) were ordered commercially [similar to Bautista et al. (2012) (NCBI accession no: NM_010290.2)]. Polymerase chain reaction (PCR) was used to amplify the Cx36 ISH probe template from big brown bat genomic DNA and add a T7 promoter. The DNA template was separated on a gel and purified. *In vitro* transcription was used to generate digoxigenin (DIG)-labeled RNA antisense probe. The probe was purified using DNase digestion and organic RNA extraction. A sense probe was also generated as a control (Fig. 1).

**FIG. 1.**
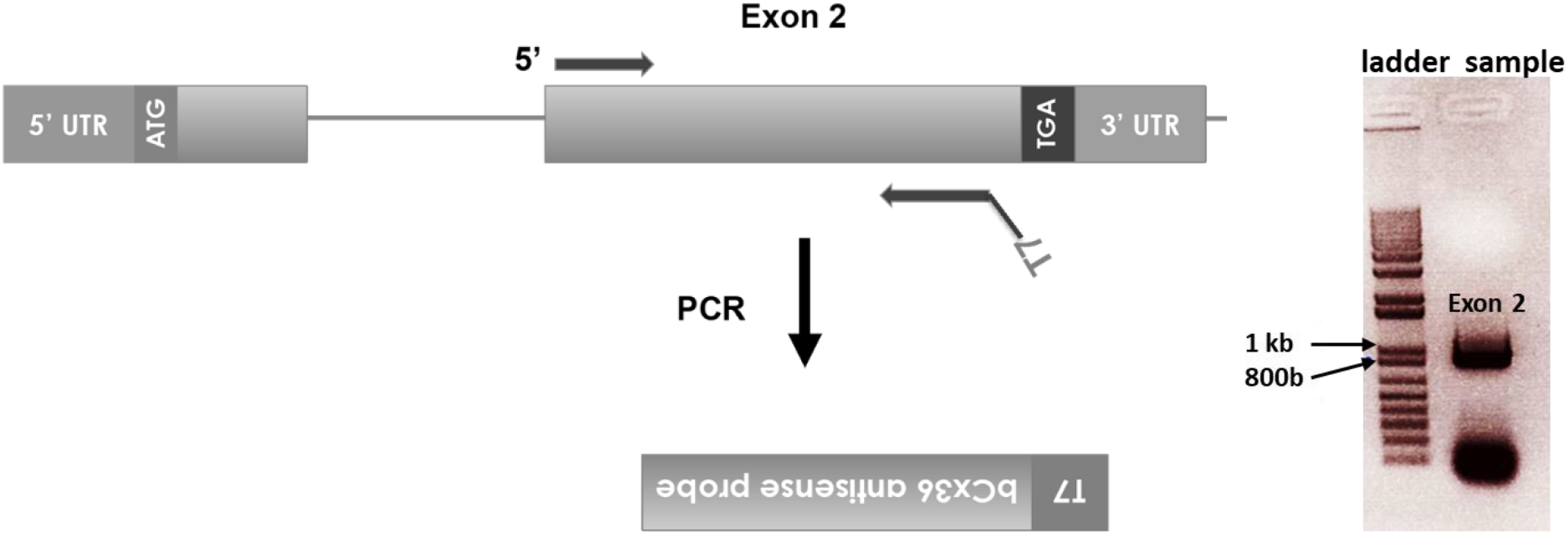
Generating DIG-labeled RNA in situ hybridization probes for Cx36 in the bat. A tBlastn search was performed using the known mouse amino acid sequence against the bat genome data and a gene that encoded a bat Cx36 ortholog was identified. Then primers were designed to amplify a sequence within the second exon that corresponded to the mouse sequence. Forward primer (5’): CCTGTTGACTGTGGTGGTGA and reverse primer (3’): TAATACGACTCACTATAGGGGCAGAATCACTGGACTGAGTCC. The bat’s genomic DNA was isolated, then custom oligonucleotides and PCR were used to amplify the probe template and to add a T7 promoter. In vitro transcription was used to generate DIG-labeled RNA sense and antisense probes.

Bats (*n* = 2) were euthanized with Beuthanasia-D, brains were removed, embedded in OCT, promptly frozen in a CO_2_ and ethanol ice bath, and then transferred to a −80 C freezer. Brains were sectioned on a cryostat, as previously described except at 10 μm thickness, using RNAse-free materials. Sections were mounted to charged slides, placed in a plastic slide box, wrapped in plastic wrap to retain moisture, and frozen at −80° C until processing. Slides were thawed (45 min to 1 hr) and sections were post-fixed to the slides for 10 min in 4% PFA. Sections were then washed in PBST, incubated twice in TEA buffer (triethanol amine with acetic anhydride) for 10 min, washed in PBST, and quenched in 1% H_2_O_2_ in PBS for 30 min. They were then incubated in Proteinase K solution for 5 min at 37° C, post-fixed, and washed. Sections were incubated in pre-hybridization buffer for 1 hr at room temperature under a hybrislip, and then hybridized with the DIG-RNA probe overnight at 65° C. Control sections were incubated in the sense probe. Sections were subjected to stringency washes in SSC (3 × 15 min at 65° C), allowed to cool to room temperature, and then washed in TBST three times for 5 min each. Sections were blocked with BSA for 30 min at room temperature, and then incubated in anti-DIG-AP antibody at room temperature in a humidified chamber. They were then washed three times for 5 min each in TBST, then incubated in detection buffer (100 mM Tris-HCl, 100 mM NaCl, and 50 mM MgCl_2_) for 5 min. Slides were incubated in 10% polyvinylalcohol with NBT/BCIP substrate and 1mM levamisole at 37° C under glass coverslips for 1 hr. Slides were then washed, post-fixed, dried, and mounted with Fluoromount-G (cat # 0100-01, Southern Biotech). Brightfield microscopy was performed on an Olympus BX60 microscope (Olympus Scientific); images were acquired using an Olympus DP72 camera and cellSens software, which was used to white-balance images. Images were then analyzed to count and measure the area of labeled cells in ImageJ by an experimenter blinded to the hypothesis using the manual Freehand tool.

#### Mouse

In order to support quantitative comparison between Cx36 expression in bat and mouse, ISH data obtained from the Allen Mouse Brain Atlas (https://mouse.brain-map.org/experiment/show/71836902) were analyzed to find percent area of expression in the dorsal and ventral CN of a male, P56, C57BL/6J mouse. The primers used to identify Gjd2 included the forward primer TGGCCATTGTAGGGGAGA and reverse primer CGAATGAGGGCAAACCTG. Detailed experimental and image capture procedures performed by the Allen Brain Institute are available in a technical white paper (Allen Institute, 2011). Images were then processed by an experimenter blinded to the hypothesis using Custom Macros (www.ijmacros.com) for ImageJ (Timothy and Forlano, 2019) in order to find the percent area (2D) of gene expression.

##### Detection of Cx36 expression pattern in heterozygous reporter mice

Mice heterozygous for Cx36/LacZ-IRES-PLAP on a C57BL/6 background (*n* = 3; Deans et al., 2001) and wild-type littermates (*n* = 1) were deeply anesthetized (0.1cc Beuthanasia) until no motor reflexes were observed, then transcardially perfused with heparinized 0.9% saline followed by fixative (1% PFA, 0.2% glutaraldehyde in PBS). Brains were dissected, post-fixed overnight, washed in PBST, serially cryoprotected with 15% and 30% (w/v) sucrose, left overnight, then embedded in OCT on dry ice and stored at −80° C. Coronal cryosections were taken at 20 μm thickness and mounted to glass slides, and then were stored in airtight containers at −80° C until further processing. After thawing, sections were post-fixed to the slides, then rinsed in PBST. Cx36 gene expression (GJD2) was revealed using a β-galactosidase assay using X-gal substrate (blue product) with nuclear fast red counterstain. Slides were washed in a staining buffer, incubated with approximately 150 μl each of 1 mg/mL X-gal (5-bromo-4-chloro-3-indolyl-β-d-galactopyranoside) in staining buffer plus 5 mM potassium ferrocyanide and 5 mM potassium ferrocyanide at 37° C until color developed. Slides were then washed in PBS for 5 min three times each, then serially dehydrated in ethanol (2 min in 50%, 75%, 90%, and 100%), then coverslipped with AquaMount and stored at room temperature. brightfield microscopy was performed on an Olympus BX60 microscope (Olympus Scientific); images were acquired using an Olympus DP72 camera and cellSens software, which was used to white-balance images. Images were then processed by an experimenter blinded to the hypothesis using Custom Macros (www.ijmacros.com) for ImageJ (Timothy and Forlano 2019) in order to find the percent area of gene expression.

## RESULTS

### Cytoarchitecture of the bat CN

The gross and cellular anatomy of the big brown bat’s CN complex have been previously described (Hall 1969; Huffman and Covey 1995; Covey 1995, 2005). Terminology and cell classification from this previous work are used here. Cell types in the AVCN have been identified as small spherical, multipolar, and globular cells (Haplea et al. 1994; Huffman and Covey 1995). Rosenberger et al. (2003) further specified spherical bushy, globular bushy, and granule cells. In the PVCN, cell types have been classified as octopus, multipolar, and elongate cells, while in the DCN, fusiform cells have been identified (Haplea et al. 1994; Huffman and Covey 1995).

The gross cytoarchitecture of the bat cochlea, 8th N, and brainstem (AVCN; IC, inferior colliculus) is illustrated in Fig. 2a. The cochlea wraps around the 8th N and the AVCN extends downward towards the cochlear spiral (Fig. 2b). The estimated total CN volume was 1.15 mm^3^, composed of the AVCN, 0.64 mm^3^, the PVCN, 0.28 mm^3^, and the DCN, 0.24 mm^3^ [not adjusted for the ^~^9.2% tissue shrinkage (Carter et al., 2004)].

**FIG. 2.**
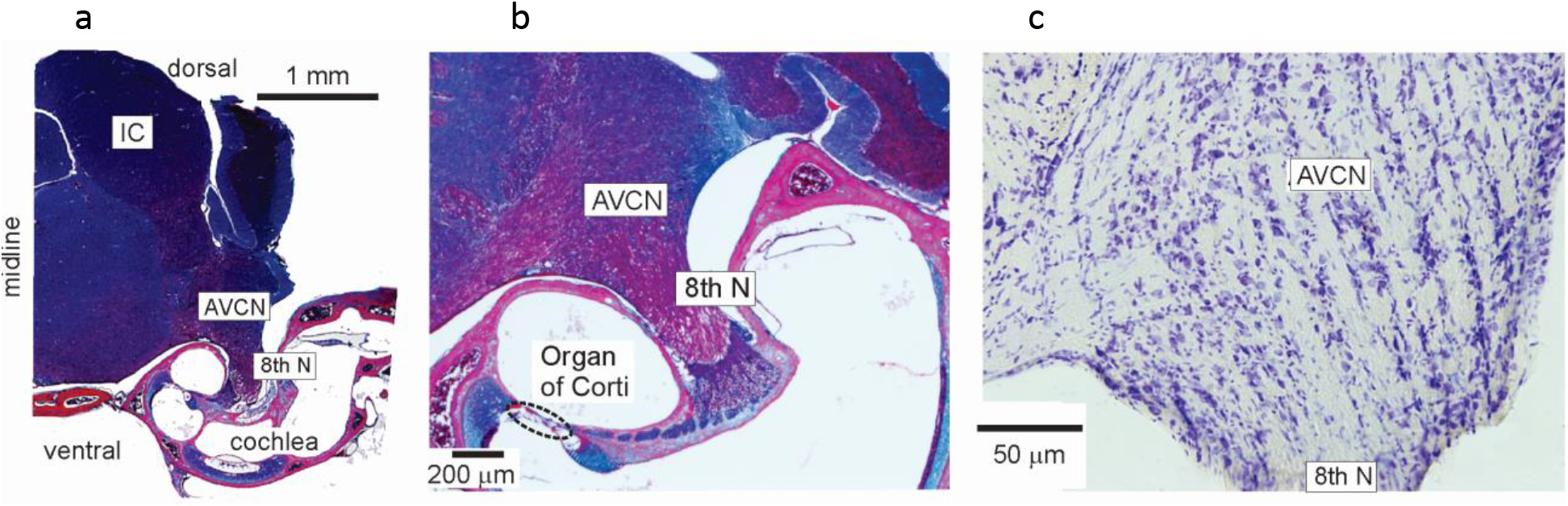
Gross morphology and cytoarchitecture of the bat brainstem, auditory nerve, and cochlear nucleus. (a) Partial coronal section taken from a whole bat head stained with trichrome, where light blue = connective tissue and decalcified bone, red = fibers, and purple/dark blue = cell bodies. (b) Closer view of (a) showing the organ of Corti (dashed black oval line), auditory nerve (8th N), and anteroventral cochlear nucleus (AVCN). (c) Entry site of 8th N into the AVCN (cresyl violet stain). Cell nuclei appear dark purple, and fibers are lightly stained, showing a slightly sloping vertical striated pattern where the 8th N is interrupted by AVCN neurons and other supporting cells.

The AVCN receives incoming auditory nerve fibers interleaved with cell nuclei in “fan-like” striations (Fig. 2c, Fig 3d; also shown in Hall 1969). Within these striations are groups of neurons (as identified using IHC-F for Map-II; Fig. 3), presumed to be globular bushy cells based on their location near the 8th N root and their size (average area = 429 ± 64 μm^2^, *n* = 6, Fig. 3g, h), round shape, and clustered cytoarchitecture (Huffman and Covey 1995; Rosenberger et al. 2003). Fig. 3d illustrates that presumed globular bushy neurons (Fig. 3c, g) are interleaved with the auditory nerve afferents (shown non-specifically labelled in green, Fig. 3b). Fig. 3e-h show a higher-magnification view of Fig. 3a-d near the entry site of the auditory nerve the ventral-most neural striation. Fig. 3h shows a total of six neurons (red) aligned in a column with a distinct cluster of three neurons with closely apposed somata (ventral portion of Fig. 3h). Bushy cell dendrites and axons are also apparent (Fig. 3c).

**FIG. 3.**
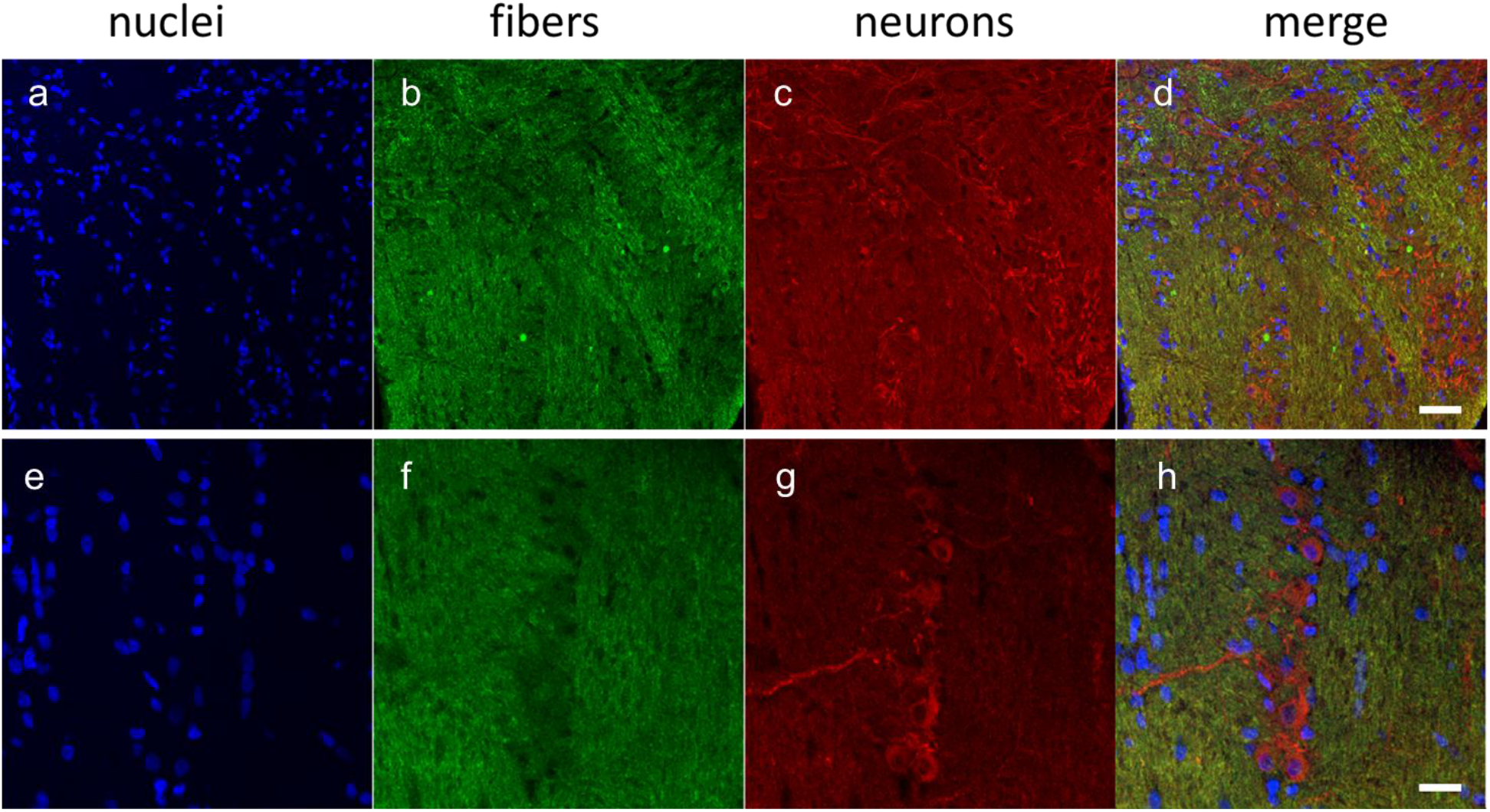
Cytoarchitecture of presumed globular bushy neurons (GBNs) in the bat’s ventral AVCN. Cell nuclei labelled with DAPI appear in blue (a & e); fibers are non-specifically labeled in green (b & f); and neurons labelled with a specific antibody to a pan-neuronal marker (Map-II) are in red (c & g). The first three panels are maximum-signal collapsed single-channel (RGB) z-stacks comprised of 5 (a-c) or 8 (e-g) individual confocal microscopy images. (e-h) a higher-magnification view of (a-d) near the entry site of the auditory nerve the ventral-most neural striation, where a column of 6 neurons is apparent, and the ventral 3 somata appear closely apposed (g, h). Scale bar = 200 μm (a-d) and 20 μm (e-h).

### Connexin36 protein expression in the bat CN

Cx36 label was visible throughout the CN. The brown DAB reaction product filled CN cells, so the subcellular location of gap junctions is not indicated with this method; however, cell bodies and their projections are readily visible. In the AVCN, Cx36 was expressed in cells whose morphology is consistent with bushy neurons (spherical and globular; mean cross sectional area = 187 ± 45 μm^2^, *n* = 11) (Fig. 4a-c). Label was also found in cells throughout the PVCN that are likely spherical bushy cells based on their round shape and location (Fig. 4g). Finally, presumed fusiform neurons in the DCN (based on location, size, and shape, mean area = 94 ± 34 μm^2^, *n* = 20, Fig. 4h, i) were labeled. Sparser label was observed in deep-layer DCN cells (Fig. 4h). No brown reaction product was observed in control sections (Fig. 4d-f, j, k).

**FIG. 4.**
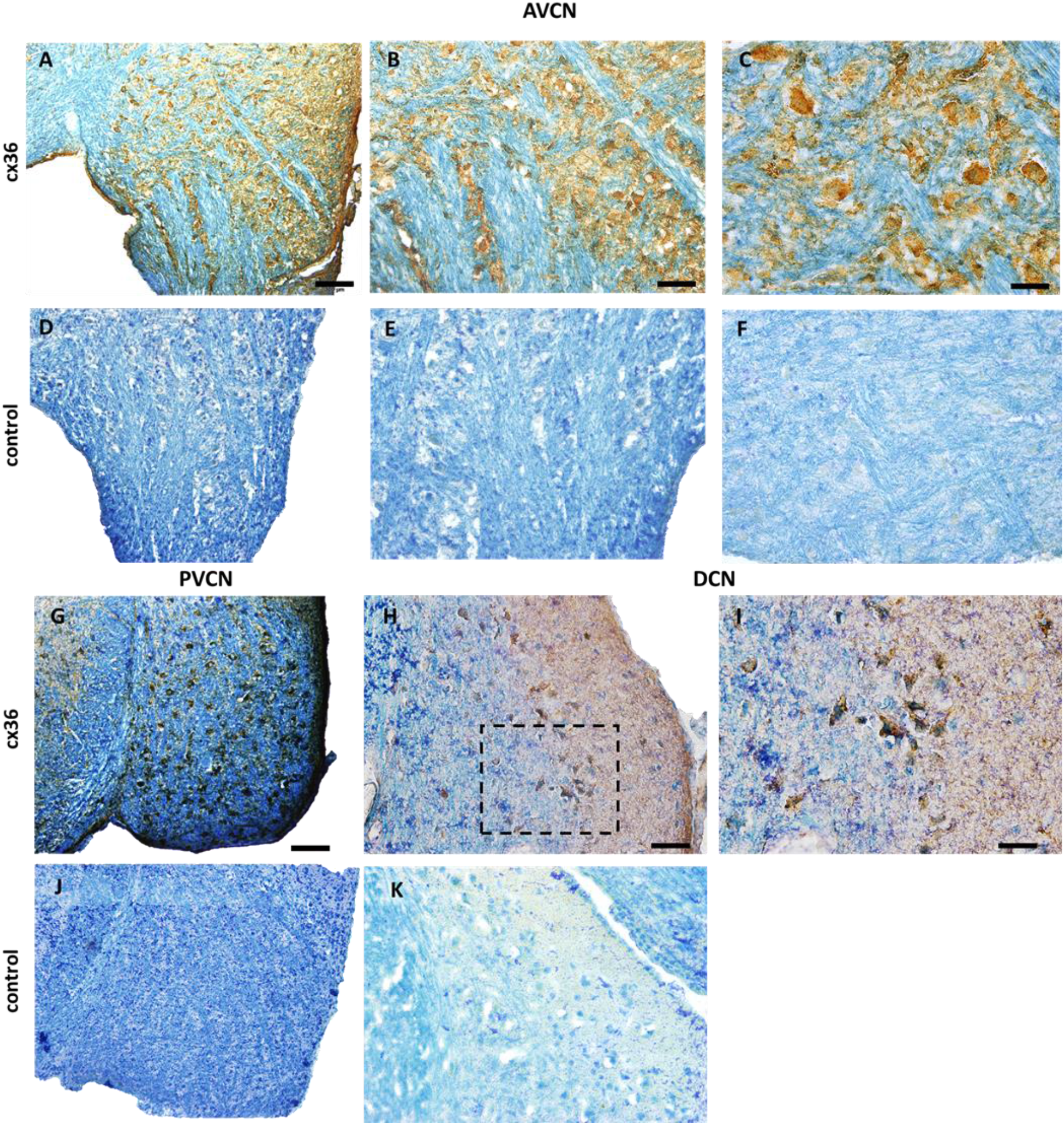
Cx36 immunohistochemistry in the bat CN. Cx36 expression in the AVCN (a, b, c). Cx36 protein was expressed throughout the AVCN near the entry site of the auditory nerve. Control samples that omitted primary antibody (d, e, f) were taken from alternating sections of the same brain. Cx36 protein was expressed heavily throughout the PVCN (g) and sparsely in the DCN (h, i). The area within the dashed box in (h) is magnified in (i), showing some Cx36-labelled cells with apparent fusiform morphology. Control sections where primary antibody was omitted showed no label (j: PVCN; k: DCN). Scale bar = 100 μm (left panels); = 50 μm (middle panels); = 20 μm (right panels).

In distinction to the IHC-DAB method that filled cells with a colored reaction product, the IHC-F method labeled characteristic Cx36 puncta (Fig. 5C, K; average area = 1.1 ± 0.8 μm^2^, *n* = 28) as well as neuronal cell bodies (Fig. 5B, J, mean area = 210 ± 37 μm^2^, *n* = 6). While subcellular localization of Cx36 cannot be definitively determined using IHC-F, puncta appeared to colocalize with neuron somata (Fig. 5c, d, k, & l), but were also visible on presumed dendrites (Fig. 5l). Cx36 was expressed in putative multipolar and octopus neurons in the ventral AVCN/PVCN marginal zone (Fig. 5c&d). Control samples that omitted primary antibody were taken from alternating sections of the same brain (Fig. 5e-h).

**FIG 5.**
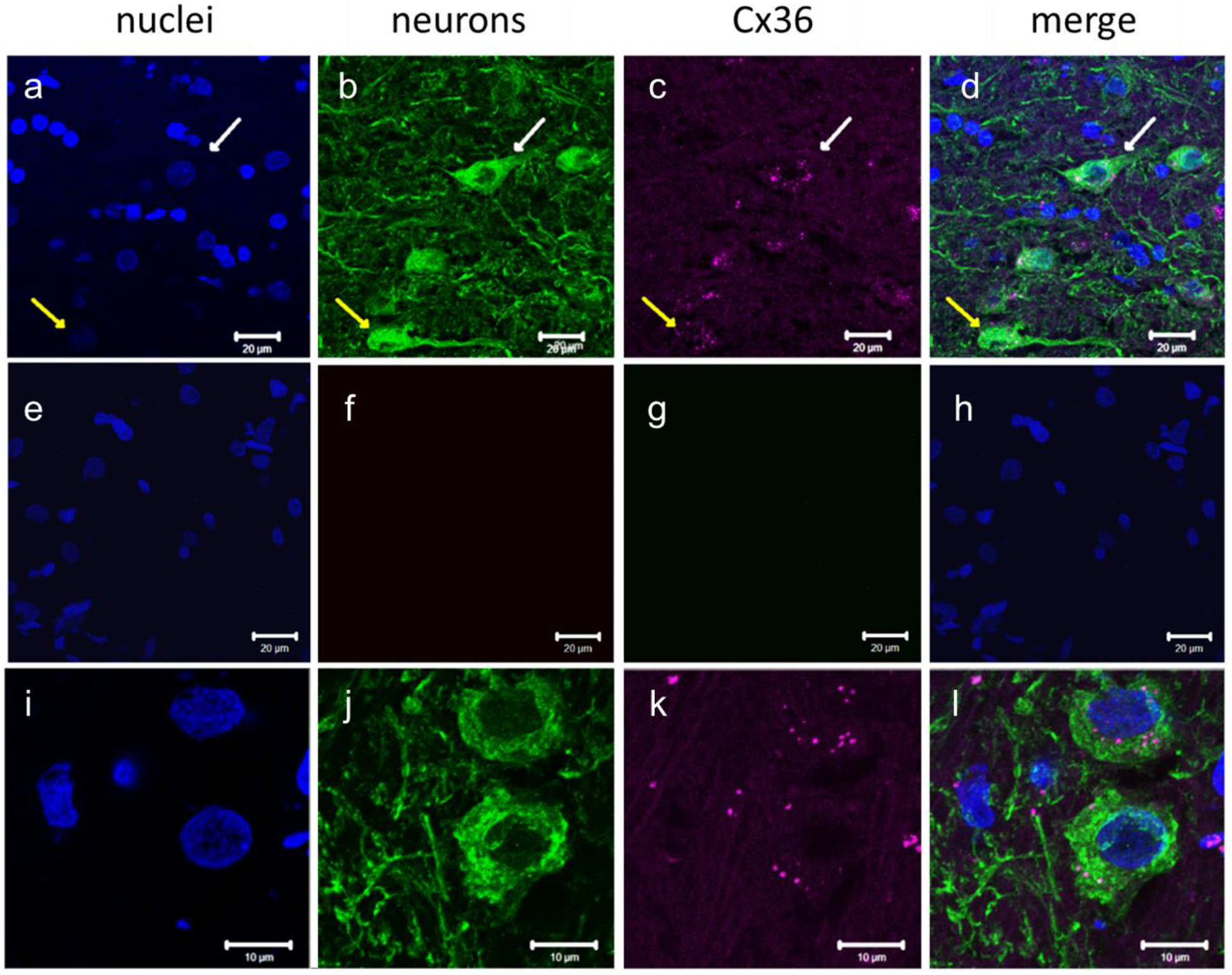
Cx36 immunofluorescence in the bat VCN. Collapsed confocal z-stack images show that Cx36 protein appears punctate (magenta, c & k) on cell bodies and dendrites of neurons in the VCN (a-d). Control samples that omitted primary antibody were taken from alternating sections of the same brain (e-h). Nuclei were stained with DAPI (blue, a, d, e, h, i, l), neurons stained with a pan-neuronal marker (green, b, d, j, l). Top row, a-d: marginal zone between AVCN and PVCN. Arrows indicate the position of two example neurons that appeared to express Cx36 puncta on their somata (white arrow = multipolar cell, yellow = octopus cell). Middle row, e-h: control sample with primary antibodies omitted. Bottom row, i-l: Nerve root area of the AVCN showing Cx36-puncta on presumed globular bushy neuron somata and dendrites.

### Genetic expression of Cx36 in the bat and mouse CN

A custom-made DIG-labelled probe was used to identify Cx36 mRNA expression in the bat CN. RNA expression was evident in the AVCN (Fig. 6a) and DCN (Fig. 6b, c, f) in the same areas where protein expression was found. Sense probe controls resulted in absence of labelling in the AVCN (Fig. 6d), and DCN and PVCN (Fig. 6e). Cx36 gene Gjd2 was expressed only in the dorsal-most portion of the posterior PVCN on the border with the DCN; sections containing the anterior PVCN could not be evaluated due to tissue quality. Gjd2 label was 9.5 ± 3% area (n = 4) and 6.8 ± 2% (n =4) area of the AVCN and DCN, respectively (Fig. 7). These differences did not reach statistical significance. In the mouse, Gjd2 was labelled in 15.8 ± 7% (n = 4) and 12.6 ± 3% (n = 4) of the dorsal and ventral CN, respectively, but differences between brain regions were not statistically significant. A Kruskal-Wallis rank sum test comparing bat and mouse data revealed that the percent area labeled for Gjd2 in the mouse samples was significantly larger than those in the bat samples (X^2^ = 6.4, *df* = 1, *p* = 0.012). Within species, no significant differences were found between the dorsal and ventral CN.

**FIG. 6.**
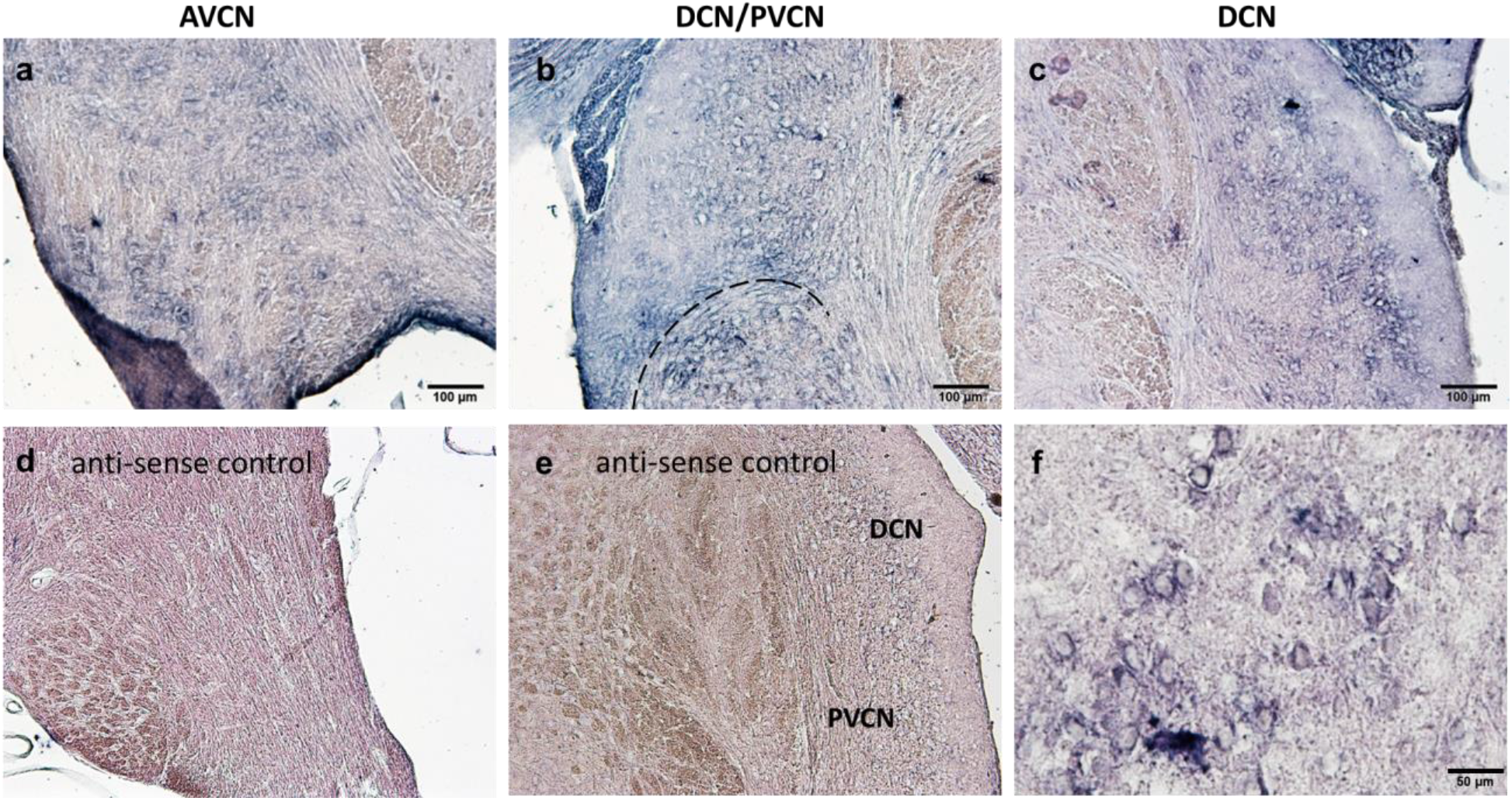
Cx36 RNA expression in the bat CN revealed by in situ hybridization. DIG-labeled Cx36 antisense (a-c, f) and sense (control, d, e) probes were applied to coronal cryosections of the bat brain. RNA expression was visualized with alkaline phosphatase enzyme digestion of NBT/BCIP, producing a purple substrate. (a) ventral AVCN, (b) DCN and dorsal PVCN separated by dotted line, (c), DCN, and (f) magnified portion of the fusiform cell layer of the DCN, showing perinuclear staining, not from the same sample as (c) or (b). Scale bar = 100 μm (a, b, d, e) and = 50 μm (f).

**FIG 7.**
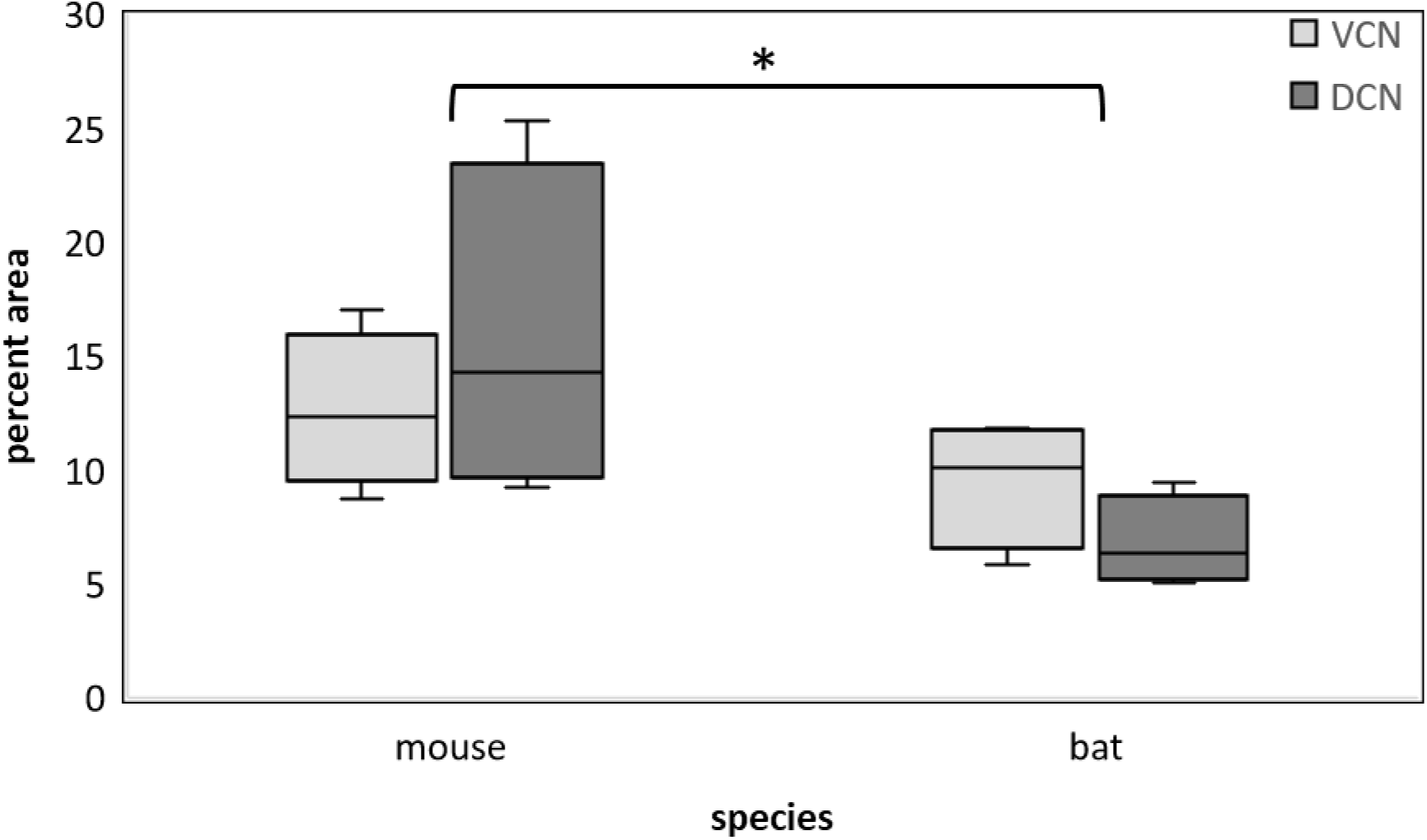
Percent area of CN labeled by Cx36 in-situ hybridization in the mouse and bat. Bat Cx36 gene Gjd2 ISH results from the AVCN (VCN) and DCN compared to mouse data from the Allen Brain Institute. Horizontal lines show the median in each box plot. VCN is light gray and DCN is dark gray. Error bars denote exclusive median upper and lower quartiles. Percent area labeled for Gjd2 in the mouse samples (VCN and DCN combined, n = 8) was significantly higher than the bat (AVCN and DCN combined, n = 8), denoted by the asterisk (*p* < 0.05).

Cx36 was expressed throughout the CN of mice with a Cx36 gene reporter (Cx36/LacZ-IRES-PLAP). Figure 8 shows the results of a β-galactosidase assay (using X-gal substrate to create a blue product), which qualitatively labelled more heavily in the DCN fusiform cell layer (Fig. 8b) and in the AVCN (Fig. 8a) than in the PVCN (Fig. 8b). Within the DCN, the deep layer also expressed Cx36, but staining was sparser than in the fusiform cell layer. In sections containing the CN, a greater total area of the CN was labelled (2.4% - 6.6% area) than other brainstem areas imaged in the same samples (0.3% - 0.8% area), which suggests a functional rather than structural role for Cx36 proteins in the CN. Label was also present where expected (in positive control regions) including the cerebellum (Fig. 8c), inferior olive, and MNTB. In the absence of the transgene that allowed cells expressing Cx36 to be labeled, wild type littermate controls showed no transgene expression.

**FIG. 8.**
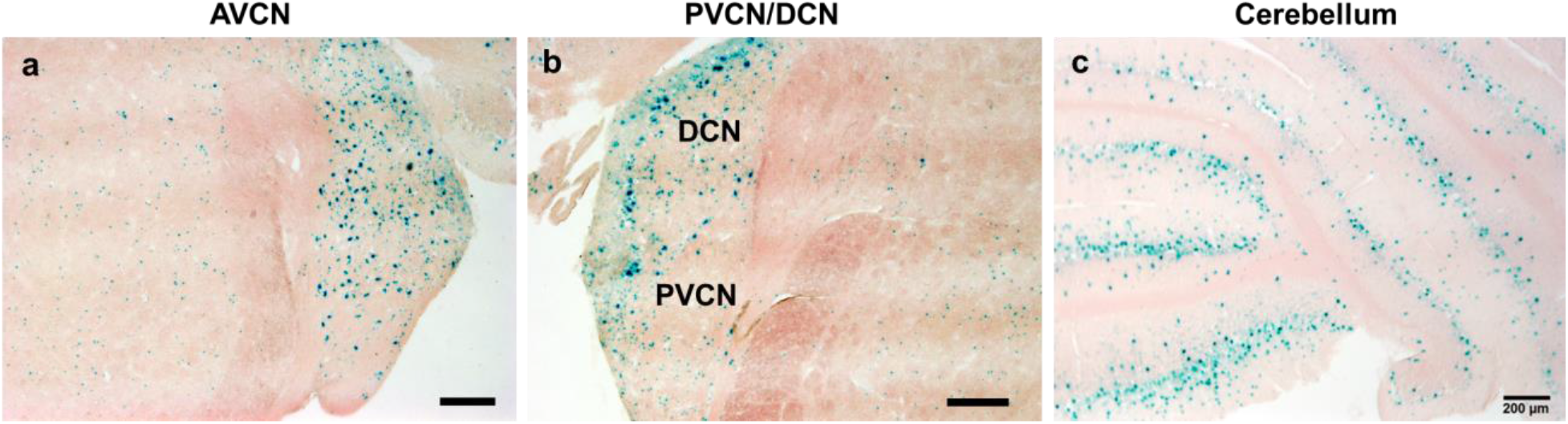
Cx36 expression in the mouse CN. GJD2 DNA (the gene for Cx36) expression was measured using a β-galactosidase assay using X-gal substrate (blue product) with nuclear fast red counterstain. Cx36 expression was abundant in the AVCN (a) and fusiform cell layer of the DCN (b). Expression was sparser in the deep layer of the DCN (b) and in the main body of the PVCN (b), but some expression was observed in the granular lamina between the DCN and PVCN (b). As expected, expression was also found in the cerebellum (c). Scale bar = 200 μm.

## DISCUSSION

Here we used two immunohistochemical labeling methods to confirm that the gap junction protein Cx36 is expressed in all three subdivisions of the big brown bat’s CN. We expanded on earlier results based on immunohistochemistry (Horowtiz et al. 2008) by measuring Cx36 expression using *in situ* hybridization to detect Cx36 RNA. In addition, in distinction to those earlier results, we demonstrated a similar distribution of Cx36 gene (Gjd2 DNA) expression in the CN of reporter mice. Qualitatively, in both the bat and mouse, there appeared to be a less robust pattern of expression in the PVCN as compared to the AVCN and DCN (Figs. 4, 8). Large multipolar and octopus neurons in the bat AVCN/PVCN marginal zone appeared to express Cx36 (Fig. 5), similar to results in the mouse PVCN described elsewhere [see Fig. 1b in Rubio and Nagy (2015)]. In addition, a larger percent area of the CN was labeled in mouse compared to bat. Our new data indicate that the anatomical substrate for Cx36-mediated electrical neurotransmission is conserved in the mammalian CN across echolocating bats and non-echolocating mice.

The differences in the methods we used to detect Cx36 expression in the big brown bat and in the transgenic mouse (i.e., mRNA target detected with alkaline phosphatase in bat vs. DNA target detected with β-gal in mouse) precludes direct quantitative comparison between Cx36 expression in the two species from the data generated in this study alone. In order to describe these differences, we analyzed publicly available in-situ hybridization (ISH) data from the mouse (Allen Institute for Brain Science, 2004). Quantitative analysis of ISH images from unique areas of the mouse VCN and DCN derived from the Allen data and of bat AVCN and DCN collected in this study showed that a significantly higher percent area was labeled for the Cx36 gene in the mouse samples. Furthermore, the average percent area for the mouse DCN was higher than the mouse VCN whereas the opposite was true for the bat, where the percent area labeled in the AVCN tended to be greater than in the DCN. We also used data from Carter et al. (2004) to estimate the volume of the bat CN. Results showed that the bat CN exhibits substantial hypertrophy (total volume 1.15 mm^3^), compared to the total volume of the mouse CN at 0.6 mm^3^ (Muniak et al. 2013, Godfrey et al. 2016). Surprisingly, a larger percent area of the mouse CN compared to the bat’s expressed Cx36. This smaller percent area of Cx36 expression in the bat (Fig. 7) suggests that Cx36 is not as concentrated or widespread in the bat as in the mouse; however, because the bat’s CN structures are hypertrophied, it is still possible that the bat’s CN contained absolutely more Cx36 expressing neurons, or that the bat’s CN contained more cells that did not express Cx36, though that was not quantitatively determined. It is unlikely but possible that 25 μm tissue sections, or debris artifact prevalent in images from the Allen atlas (see https://mouse.brain-map.org/experiment/show/71836902) could have accounted for the higher percent area measured in the mouse. In any case, the non-statistically significant trend that the bat expressed Cx36 in a larger percent area of the AVCN compared to the DCN deviated from the mouse results (Fig. 7) and might indicate that the bat’s hypertrophied AVCN may contain more neural circuits for electrical coupling than in the mouse.

In a previous study based on IHC-F, Horowitz et al. (2008) observed Cx36 expression in dorsal regions of the big brown bat’s AVCN, with sparse and uneven label in the PVCN, and no label in the DCN. Our results confirm robust expression throughout the bat AVCN, but, in contradiction to these earlier results, we found expression in the DCN, especially in the fusiform cell layer. We also observed expression in marginal zone areas where the PVCN meets the ACVN and DCN (Figs. 4g, 5, 6b), and in the nerve root area of the AVCN (Figs 4a and 6a). The bat’s anterior PVCN showed more evenly distributed label for Cx36 using IHC (Fig. 4g), but the posterior PVCN showed more label near the dorsal margin with the DCN (Fig. 6b). There are several possible reasons for the differences between Cx36 expression reported by Horowitz et al. (2008) and the present study. First, the previous work relied on thicker, 50 μm-thick brain sections while thinner, ≤ 20 μm cryosections were used here. Second, that study’s methods did not include an antigen retrieval step. Here, an antigen retrieval step was used to enhance Cx36 detection, as 4% paraformaldehyde might result in over-fixation. Both of these differences could have contributed to less-than-optimal penetration of the antibody in the 2008 study (for a detailed discussion, see Rubio and Nagy, 2015). Since the present study was conducted prior to 2015, additional improvements to tissue fixation methods as suggested by Rubio and Nagy (2015) were not implemented here. Likely for similar reasons, Horowitz et al. (2008) also did not observe Cx36 expression in the CN of mice (strain unknown). Here, we did not use immunohistochemistry to identify Cx36 expression in the mouse – rather, transgenic mice allowed for an assay to be used which was less sensitive to fixation. We found qualitatively less expression in the mouse PVCN (Fig. 8b) as compared to the bat (Fig. 4g). Robust Cx36 expression was detected in the fusiform cell layer of the DCN, and some label was found in all layers (see Fig. 4b & 6c for bat; Fig 8b for mouse).

The location of Cx36 expression within the big brown bat’s DCN (Fig. 4h) suggests that electrical transmission may also occur in DCN neuron types (such as Giant cells) other than fusiform, stellate, and Golgi cells, which have been characterized previously in other mammals (Apostolides and Trussell 2013; Apostolides and Trussell 2014a, b; Yaeger and Trussell 2016). Of all CN subregions, the functional auditory circuitry in the DCN is the best understood, and research has shown that sub-threshold principal neuron (fusiform cell) activity drives inhibitory stellate neuron firing with precise timing through electrical synapses (Apostolides and Trussell 2014a). Additionally, Golgi cells predominantly form electrical synapses with one another, creating a complex network that mediates both excitation and inhibition (Yaeger and Trussell 2016). There are little electrophysiological data from FM bats that address the relationship between excitation and inhibition in DCN circuitry (Suga 1964; Haplea et al. 1994; Covey and Casseday 1995; Covey 2005).

The results from previous work combined with our study suggest that electrical neurotransmission may also play a primary role in the bushy cell network in the AVCN in bats as in other species (Gomez-Nieto and Rubio 2009, 2011). The pattern of expression in the bat AVCN appears similar to that in the mouse, notwithstanding the differences in percent area expression (Fig. 7) and the hypertrophy of the bat CN (Covey 2005). In the bat, groups of bushy neurons expressing Cx36, likely coupled by gap junctions, were interleaved with auditory nerve fibers, forming prominent striations (Figs. 3, 4a, 4b, 5). These striations were not apparent in the mouse using the β-galactosidase assay and nuclear fast red counterstain (Fig. 7a). The functional significance of these organizational differences is unknown. The pattern of CX36 expression in the bat’s ventral AVCN resembled that in the rat, in that grouped Cx36-expressing AVCN neurons were located distally in the 8th nerve root in both species (see Figs. 3, 4, & 7) (Gomez-Nieto and Rubio 2009; Rubio and Nagy 2015). In the mouse and rat, Rubio and Nagy (2015) found mostly pairs of bushy cells coupled by Cx36 puncta, and occasionally found 3-4 cells linked in sequence by puncta. Networks of 5-6 bushy cells have been described in the rat and rhesus monkey AVCN, where they are coupled by diverse gap junctions, receive primary excitatory inputs, and inhibition (Gomez-Nieto and Rubio 2009, 2011). Similar cytoarchitecture was observed in the bat in this study (Fig 3). Perhaps the most interesting feature of globular bushy neurons, demonstrated in the rat by Sotelo et al. (1976) and Rubio and Nagy (2015), is that they receive mixed electrical and chemical primary auditory inputs, and also exhibit purely electrical connections with one another [mediated by Cx36 (Rubio and Nagy 2015)]. In the mouse, networks of bushy cells in the rostral AVCN receive more non-primary inhibitory and excitatory inputs than in the caudal AVCN, where the majority of primary auditory nerve fibers terminate (Lauer et al. 2013). Those findings and the data presented here suggest that Cx36 is expressed at mixed primary auditory afferent-bushy cell synapses and inhibitory interneurons of the bat’s AVCN.

Functionally, electrical coupling between bushy cells in the AVCN might underlie the enhanced temporal synchronization of AVCN output (Rubio 2018), particularly for comparing the timing of spikes in different, closely spaced frequency channels (Rosenberger et al. 2003; Covey and Casseday 1999; Covey 2005; Curti et al. 2012; Pereda 2014; Rubio and Nagy 2015; Nagy et al. 2019). This could be the early auditory computational basis for accurately detecting coincident inputs, which is essential for two features of echolocation: 1) harmonic component analysis and 2) echo delay processing (Simmons et al. 2003; Bates and Simmons 2011; Simmons 2014; Finneran et al. 2020). It is possible that gap junctions in neuronal circuitry of the CN enhance temporal registration of echo waveforms relative to what would occur from solely chemical synapses. However, and in contrast to the conclusions of Horowitz et al. (2008), the results presented here suggest that the presence of Cx36 in the big brown bat’s CN complex alone does not reflect unique neural mechanisms related to echolocation. Instead, the prevalence of Cx36 in mouse CN indicates that synaptic mechanisms for enhanced temporal acuity in echolocation might be more difficult to observe, for example, the number or size of Cx36 puncta or the origin and nature (electrical, chemical, or mixed) of auditory afferent synapses on a given bushy cell, or the spatial arrangement of bushy cell clusters. The finding that in the bat a larger percent area of the AVCN than the DCN expresses Cx36 suggests a functional difference between AVCN and DCN circuitry in temporal processing. There is little known, however, about the operation of the CN complex in FM bats (Suga 1964; Haplea et al. 1994); even in constant-frequency bats (such as the mustached bat, *Pteronotus parnellii*; Marsh et al. 2006) with their different mode of echolocation, our understanding of sound processing in the CN remains limited. Comparisons between AVCN auditory afferent synapses and inhibitory interneuron synapses onto bushy cells in echolocating bats and non-echolocating species, especially those like the mouse that hear ultrasound, could provide further insight into fundamental auditory brainstem processing mechanisms and specializations for echolocation.

## ACKNOWLEDGEMENTS

We thank Gilad Barnea for providing mouse specimens, Carolina Veltri for assistance in data collection, and Kelsey N. Hom for ImageJ software advice. Portions of this work were presented at the meetings of the Acoustical Society of America (2012), Society for Neuroscience (2012), and the Association for Research in Otolaryngology (2014). This research was supported by the Office of Naval Research (N00014-14-1-05880) to JAS, and by an Office of Naval Research Multidisciplinary University Research Initiative (N00014-17-1-2736) to JAS and AMS.

